# Unveiling BPDE Induced Carcinogenic Signaling: Computational Insights into NF-κB, MAPK, and PI3K/Akt Pathway Activation

**DOI:** 10.1101/2025.01.08.631754

**Authors:** Karishma Vivek Kathpalia, Anil Kumar, Awadhesh Kumar Verma

**Author notes:** Corresponding author: Awadhesh Kumar Verma.

## Abstract

The Carcinogen benzo[a]pyran-7, 8 dihydrodiol 9, 10 epoxide(BaP-DNP) the active metabolite of benzo[a]pyrene has been postulated to induce carcinogenesis through formation of adduct with DNA and through alteration of critical pathways involved in signaling. Besides genotoxicity, BPDE has more recently been reported to alter the NF-κB, MAPK, and PI3K/Akt, that are involved in inflammation, cell survival and cell proliferation. From the above molecular docking analysis of BPDE binding to the potential proteins of these pathways, the mechanism of action of the compound is examined. The studies conducted on docking reveal moderate binding affinity of BPDE with NF-κB, that is, -6.11 kcal/mol, which points out its role in inflammation and oncogenic signaling. In addition, BPDE has more protein-binding affinities within the MAPK and PI3K/Akt pathway for CDK1 (-7.46 kcal/mol), LOX (-7.47 kcal/mol), and CDK6 (-6.84 kcal/mol). These indicate that influence of BPDE in tumour progression should not be only through the activation of NF-κB but would rather be supported by significant inputs of pathways that include PI3K/Akt and MAPK. This interaction would seem to favor cell survival, proliferation, and apoptosis resistance. Broader analysis of the data presented allows defining NF-κB as one of the key molecules contributing to BPDE-induced carcinogenesis; however, the data also point to the significant participation of the MAPK and PI3K/Akt pathways in this process. Future work should also more fully detail how these pathways interconnect and also evaluate the effectiveness of inhibiting NF-κB and related pathways in preventing BPDE’s carcinogenic action. This study adds to understanding of molecular mechanisms and therapeutic targets of BPDE.

**Graphical Abstract:** 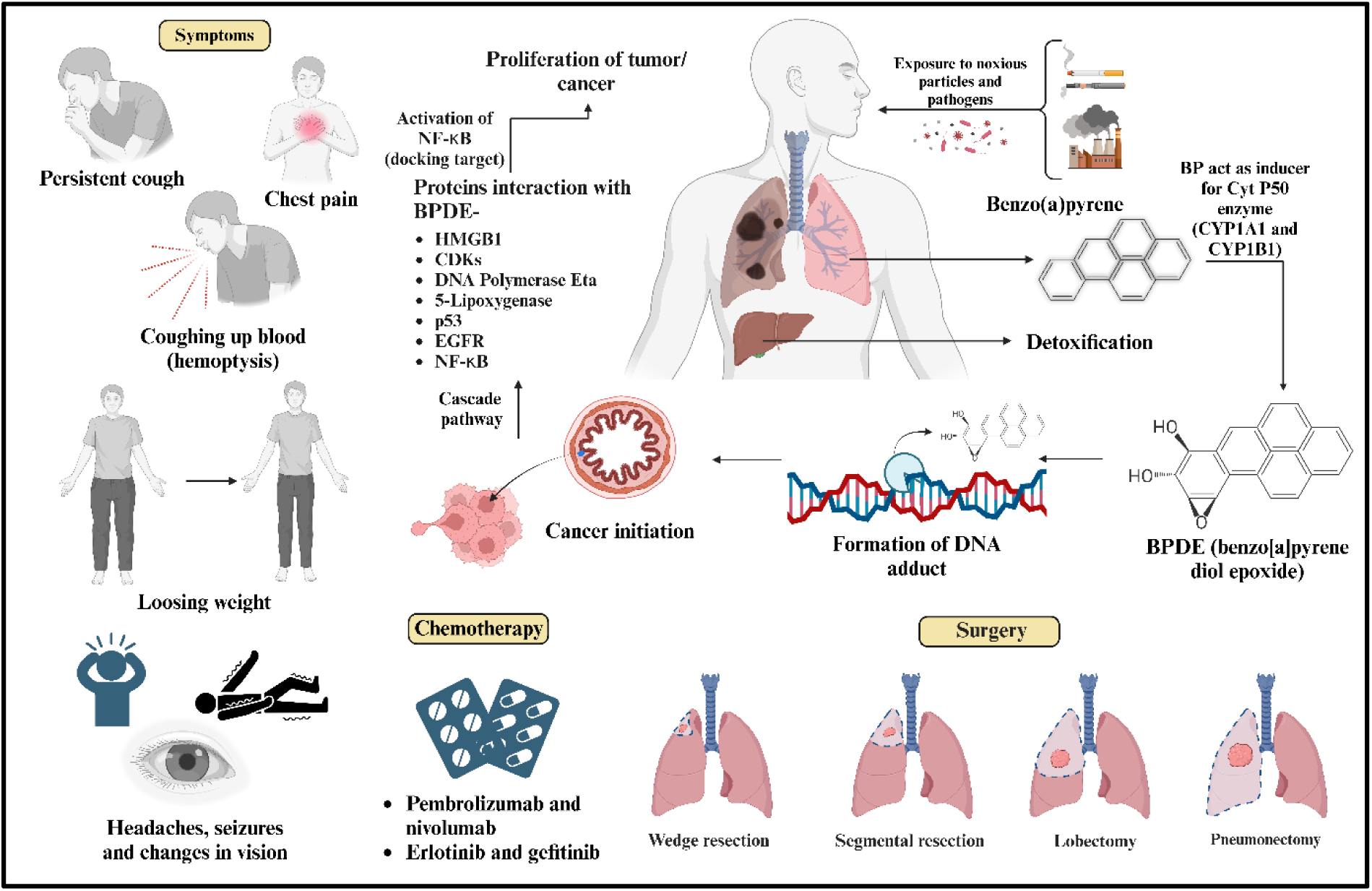

## 1. Introduction

For effective evaluation of the carcinogenic potential of BPDE, understanding of the interactions of this product with key signaling pathways is essential. BPDE is known as one of the metabolites from the polycyclic aromatic hydrocarbon Benzo[a]Pyrene (BaP), representing a powerful environmental carcinogen that induces DNA damage, thus activating a variety of oncogenic signaling pathways: NF-κB, MAPK, and PI3K/Akt1–3^1–3^. Stimulation of these pathways is imperative because they’re related to altered cell signals, inflammation, and tumour formation, causing lung cancer. First to last stage symptoms for lung cancer are persistent cough, headache, chest pain, coughing up blood, loossing weight, change in vision and seizure. BPDE Interaction with NF-κB, a transcription factor: when activated, translocated to the nucleus, activates the expression of inflammation response and survival genes^3,4^. BPDE Causes phosphorylation of IκBs, leading to the disappearance of IκBs while NF-κB is translocated to the nucleus where it is released^5^. This is a critical step for the transcription of pro-inflammatory cytokines, like IL-6, which further supports the inflammatory microenvironment that fosters cancer development. The binding affinity of BPDE to NF-κB, implies that BPDE can directly impact NF-κB activity, thereby enhancing its transcriptional potential^6^.

BaP is one of the PAHs that is prevalent in cigarettes smoke. Once inhaled, BaP is metabolized in the lungs to different active metabolites by the enzymes CYP1A1 and CYP1B1^14,15^. This metabolic activation of BaP results into a formation of a highly reactive molecule BPDE. Interaction of BPDE with various molecules ultimately leading to cell proliferation and growth. BPDE is important in the carcinogenic process as it can cause DNA damage and possibly subsequent mutations^16^. The principal BPDE/DNA interaction involves covalent binding at guanine bases of DNA, which is the well-known ultimate stage of carcinogenesis. BPDE accumulates at the N2 position to form bulky adducts, thus disrupting base pairing and DNA replication^17,18^. Its presence frequently leads to G to T transversions during DNA replication when thymine lies opposite the modified guanine base^19^. Such mutations disrupt genomic stability and cellular functions. Apart from direct DNA damage, BPDE triggers epigenetic changes, specifically histone modification and DNA methylation^20^ . BPDE reduces the histone acetylation level that results in a compact chromatin structure, lowers the gene expression level, and hampers the transcription of cell cycle and apoptosis regulator genes^21,22^. It has also been documented that BPDE enhances the hypermethylation of cytosine in the gene promoter regions in the CpG islands, hence suppressing the action of tumour suppressor genes whose functions are for cellular homeostasis and preventing uncontrolled growth^23^. The BPDE changes the microRNAs, miRNAs involved in the post-transcriptional regulation of genes. It downregulates tumour suppressor miRNAs, including let-7 and miR-34, that regulate oncogenes by inhibiting cell cycle arrest^24^. These activities result in an increased amplitude of oncogene activation and downregulation of tumour suppressor genes that increase the rate of cellular proliferation and survival. The complex interaction between genetic alterations induced by BPDE and epigenetic modifications generates a genomic instability cell transformation cycle. These confer a growth advantage that drives cancer initiation and progression into malignancies^25^ Figure 1 (a).

**Figure 1.**
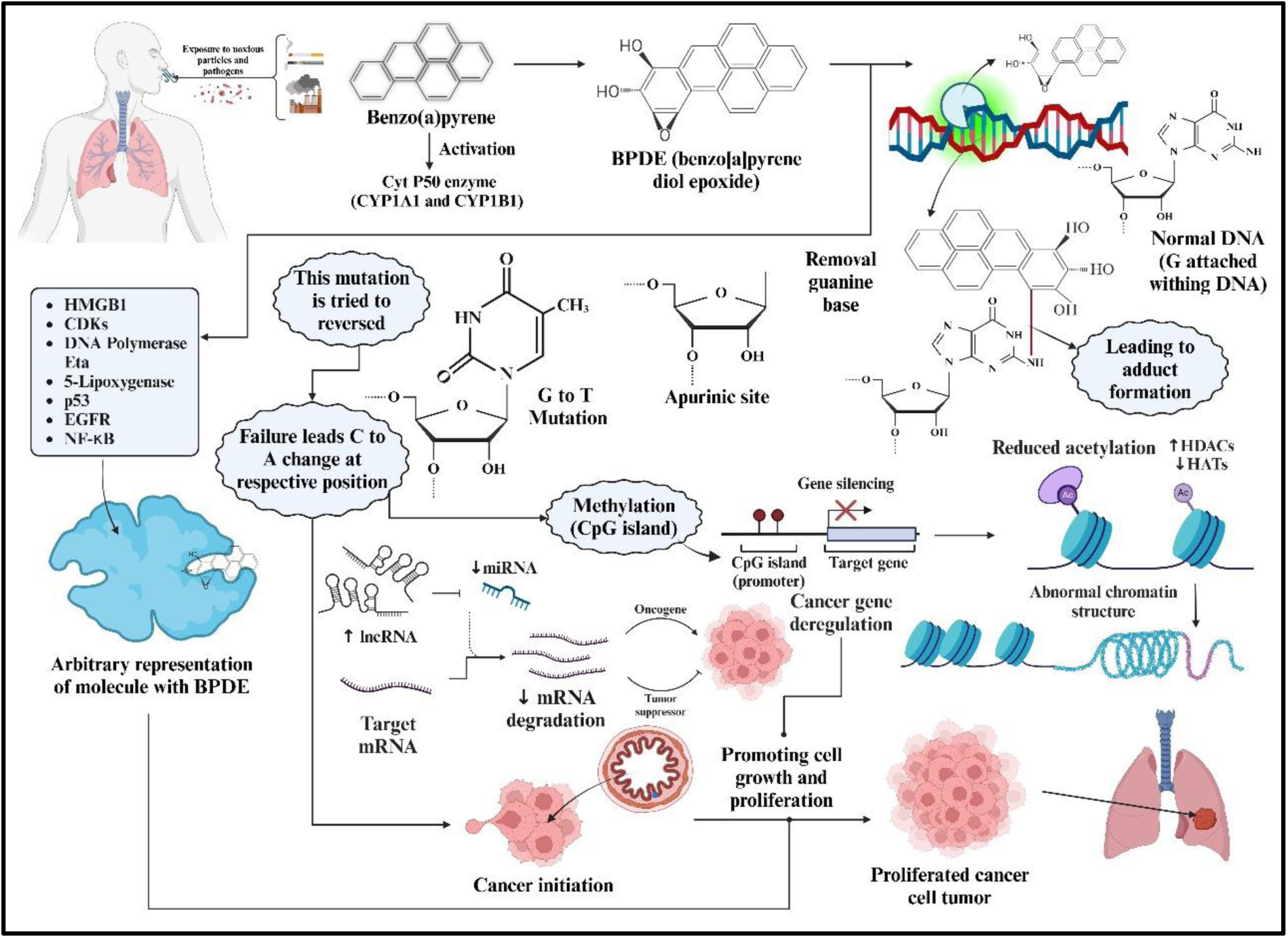
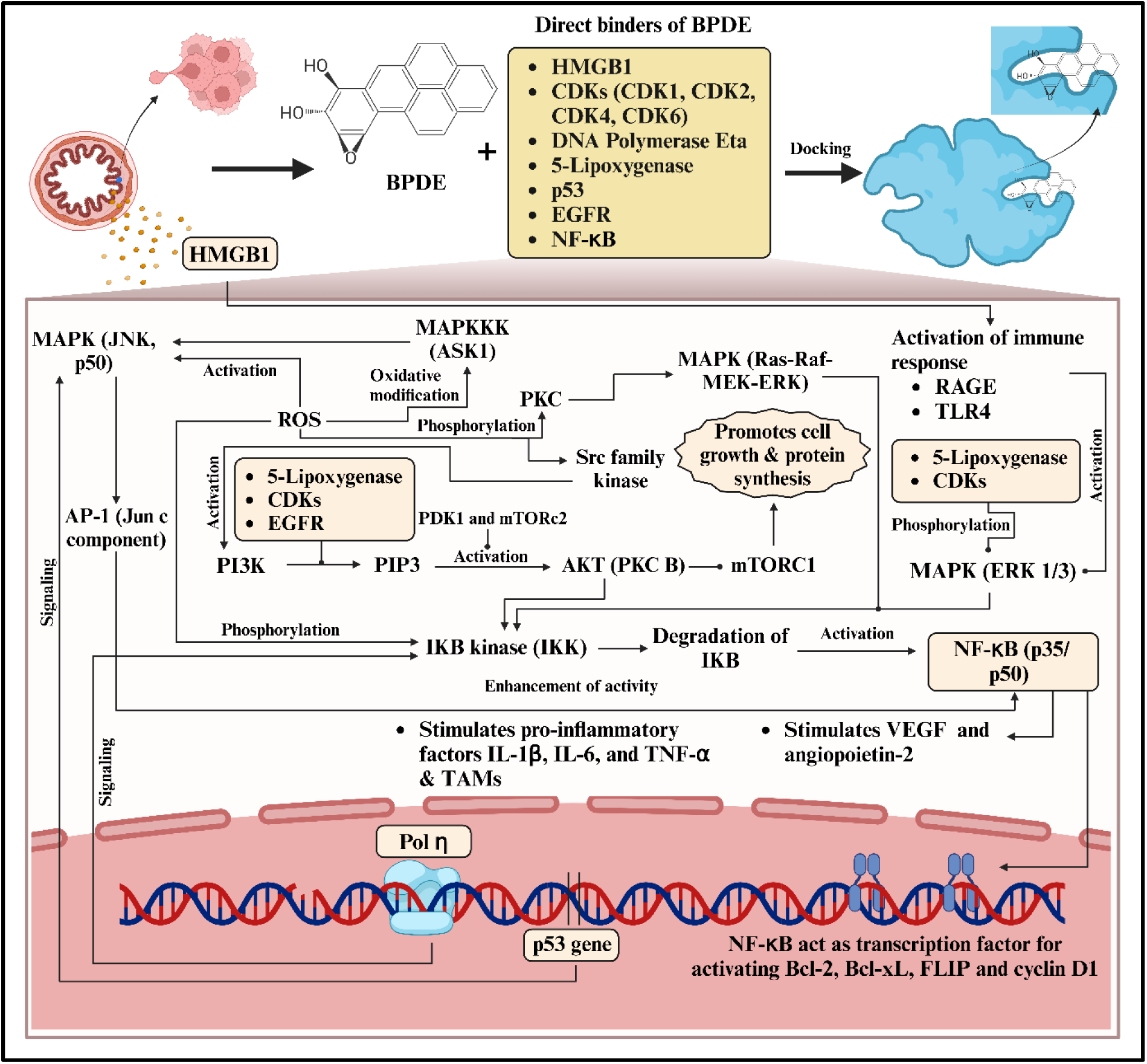
(a) It depicts the genetic level changes that take place from the point BPDE enters the body to till the cancer cell proliferation changes (b) Molecular cascade mechanisms of BPDE- induced NF-κB activation and subsequent signaling pathways

The illustration in Figure 1 (b) discusses the molecular mechanisms activated by the BPDE, a reactive BaP metabolite, primarily regarding its docking interactions and how it impacts downstream effects upon key signaling pathways, with a particular focus on the NF-κB, MAPK, and PI3K/Akt pathways. BPDE directly interacts with several critical biomolecules, such as HMGB1, cyclin-dependent kinases (CDKs), p53, Pol η, 5-lipoxygenase, EGFR, and NF-κB. These interactions initiate a cascade of signaling events that significantly impact cellular processes^4,26^. The MAPK signaling pathway is activated through MAPK (JNK, p50) and MAPKKK (ASK1), driven by oxidative modifications. This activation triggers downstream signaling, including the activation of AP-1 (Jun c component), which regulates transcriptional responses associated with inflammation and cellular stress^9^. Altogether, the implementation of the pathway of PI3K/Akt also occurs, when LOX, CDKs and EGFR leading to the phosphorylation of PI3K through BPDE forms PIP3 which activates AKT through the action of PDK1 and, enhancing survival potential, protein synthesis hence enabling cellular stress resilience^26^. In NF-κB signaling pathway, when activates the IKK, BPDE leads degradation of IκB and consecutive releases of NF-κB (p35/p50). NF-κB can be described as a transcription factor that in turn regulates the expression of anti-apoptotic genes including Bcl-2, Bcl-xL, and Inflammatory cytokines such as IL-1β, IL-6 TNF-α, respectively^26,27^. All of these pathways together facilitate inflammation, immune activation and cell survival. BPDE activates immune response pathways through receptors such as RAGE and TLR4, which recognize BPDE-modified proteins and amplify inflammatory signaling^28^. This activation process also includes the participation of ERK1/3, which phosphorylates NF-κB targets, further increasing the transcriptional activity^29,30^. BPDE promotes tumour growth and metastasis by initiating angiogenic factors, such as VEGF and angiopoietin-2^26^. It also binds p53, CDKs and activates diverse inflammatory mediators including IL-1ß, IL-6, TNF-α and TAMPs, leading to control of cell cycle mechanisms that triggers carcinogenicity^4^. In addition to the direct activation of NF-κB BPDE also activates MAPK and PI3K/Akt signaling pathways. Such signaling molecules interact in a network of signaling pathways that not only promotes the conversion of normal cells to cancerous ones and the maintenance of the inflammatory state characteristic of tumour environments^30,31^. Studies have demonstrated that BPDE can form DNA adducts, through carbon atom of epoxide ring and nitrogen atom of guanine mainly at N2 position, leading to mutations and genomic instability^32^. Due to BPDE induced DNA damage, exposure biomarkers pave way for early detection and possible treatment^33,34^. Information about molecular events associated with activation of NF-κB and other signaling pathways by BPDE not only advances understanding of the pathophysiology of BPDE-induced carcinogenicity, but also identifies targets for intervention that could attenuate the effects of this environmental carcinoma. Interaction with cytoplasmic NF-κB and subsequent enhancement of the activated protein and MAPK and PI3K/Akt pathways provide additional support for a multidimensional approach in understanding the BPDE carcinogenic process. For each of the mentioned molecules for docking, their BPDE interaction and role in cell proliferation sum – up from Figure 1(b) can be gathered from Table 1.

**Table 1.** Molecule specifications which are affected by BPDE directly induced causing cancer proliferation and survival.

| **Molecule** | **BPDE Interaction** | **Role in Cell Proliferation** | **Pathway Involved** | **References** |
| --- | --- | --- | --- | --- |
| **HMGB1** | BPDE binds directly to HMGB1, affecting its role in DNA repair and inflammatory signaling. | Regulates DNA repair and inflammatory response. | MAPK | 7 |
| **CDK1, CDK2, CDK4, CDK6** | BPDE interferes with kinase activity, causing abnormal cell cycle progression. | Regulate cell cycle checkpoints for cell proliferation<sup>8</sup>n. | PI3K/Akt and MAPK | 9 |
| **DNA Polymerase Eta (Pol η)** | BPDE-DNA adducts block polymerase activity, leading to replication stress. | Involved in bypass replication to repair damaged DNA. | MAPK | 8 |
| **5-Lipoxygenase** | BPDE modulates enzyme activity, disrupting inflammatory and proliferative signals. | Plays a role in inflammatory mediator synthesis and cancer progression. | PI3K/Akt | 10 |
| **p53** | BPDE causes p53 mutation, impairing Tumour-suppressing activities. | Tumour suppressor regulating cell cycle arrest, apoptosis, and DNA repair. | MAPK | 11 |
| **EGFR** | BPDE activates EGFR, promoting downstream signaling cascades. | Drives cell survival, growth, and migration via receptor tyrosine kinases (RTK). | PI3K/Akt and MAPK | 12 |
| **NF-κB** | BPDE triggers signaling cascades, ultimately leading to NF-κB activation. | Transcription factor that regulates immune response, inflammation, and cell survival. | NF-κB | <sup>13</sup> |

### 1.1. Research gap

The mechanisms by which BPDE brings about carcinogenesis are still not well understood; this hampers the formulation of accurate therapeutic plans. The existing drugs that target EGFR include Gefitinib (Iressa), Erlotinib (Tarceva), Afatinib (Gilotrif), Dacomitinib (Vizimpro), Osimertinib (Tagrisso), and the latest Amivantamab (Rybrevant). These drugs are primarily effective in the early stages of cancer, but their efficacy is significantly limited in later stages due to the development of drug resistance^35^. Surgical measures can occasionally be the best solution if the disease is already in the advanced stage, with high cost^36^. This work explored whether BPDE activates NF-κB independently of the activation of MAPK and PI3K/Akt. By analyzing BIOMPC’s level 2 putative molecular targets, BPDE interacts with key molecules including HMGB1, CDKs (1, 2, 4, and 6), p53, DNA Polymerase Eta, and 5-LOX^37,38^. By unraveling these interactions, the study intends to identify new avenues for targeted treatment and distinct therapeutic modalities that could help in designing drugs with enhanced efficacy to prevent and treat drug-resistant advanced and refractory cancers.

## 2. METHODOLOGY

### 2.1. *In silico* molecular modeling and optimization of BPDE as ligand molecules

3D structure of BPDE was modeled through Avogadro software. Every time 2D and 3D cleaning was performed and 3D structural confirmation of the molecule was checked by visualizing them in Avogadro view. Further, the calculation was done for each and every parameter like atomic coordinates (XYZ), bond length, and bond angle using ChemDraw software and optimized the structure up to the transition state followed by energy minimization using molecular mechanics 2 (MM2) at intermediate level using Chemdraw and Chem3DPro. Final optimization of BPDE was performed via Density Functional Theory (DFT) approach using Gaussian 09 software package. RB3LYP functionals with 6-311G basis set were used in combination for ensuring the high level of accuracy^39–41^. The optimized ligand was further used for molecular docking analysis to find interaction of receptor-ligand complex.

### 2.2. Preparation of Receptor molecules

Protein structures of NF-κB with PDB ID: 1NFI, HMGB1 with PDB ID: 2HDZ, CDKs with PDB IDs: 4YC6, 3PXR, 3G33, and 4AUA, Pol η with PDB ID: 1JIH, 5-Lipoxygenase with PDB ID: 7TTK, p53 with PDB ID: 2OCJ, and EGFR with PDB ID: 3VJO were downloaded from the RCSB Protein Data Bank to be used for docking study, in order to analyze the binding affinities and energies of BPDE conjugates. The NF-κB structure, which is derived from Homo sapiens and expressed in Escherichia coli, has a resolution of 2.70 Å, with a dimer (A & C chains) of 301 amino acids, and was prepared by cleaning crystallographic water molecules and adding polar hydrogens, Kollmann charges, and Gasteiger charges. HMGB1, also from Homo sapiens, has a resolution of 2.00 Å, with 91 amino acids in a single A chain. CDK1 and CDK2 belong to the one protein with the resolutions of 2.60 Å and 2.00 Å, respectively, whereas CDK4 and CDK6 belong to the one protein with the resolution of 3.00 Å and 2.31 Å; the docking studies were done for each conjugate separately and each of the protein contains five chains A, C, E or G with different amino acid counts. Pol η is derived from Saccharomyces cerevisiae and consists of 513 amino acids and a dimer resolution of 2.25 Å For, 5- Lipoxygenase, derived from Homo sapiens comprises of 691 amino acids and a dimer resolution of 1.98 Å. Homo sapiens shared p53 structure on tetrameric form A, B, C, D chains and 219 amino acids which has a resolution of 2.05 Å. Finally, EGFR, expressed in Spodoptera frugiperda, has a dimeric structure with 334 amino acids and a resolution of 2.64 Å. In each case, docking was performed to find the binding affinity and energy of the BPDE conjugates, with the use of Discovery Studio and PyMol for visualization and analysis.

### 2.3. Molecular Docking

Rigid molecular docking was performed with the optimized protein molecule using Autodock 4.2 tools. Genetic algorithm simulation program was used with a population size of 300 and 100 runs. The remaining parameters were kept as it is to their default values. The output docked file was saved as Lamarckian GA. Top 10 confirmations of protein-ligand complex were saved based on their negative binding energies (ΔG) and RMSD values. PyMol and Discovery Studio software were used to study the interaction and binding energy of conjugates.

## 3. Results and Discussion

### 3.1. *In silico* approach for molecular modeling and docking of BPDE-TNF-α complex

Figure 2 shows the optimized configuration of BPDE in ball and stick model generated using Gaussian09 software through DFT approach. The grey Colour ball represents the C atoms, white Colour ball as hydrogen atoms and red Colour as oxygen atoms in configuration of BPDE. The entire structure has four benzene rings and one hexagonal structure attached with oxygen atoms (represented through red colour).

**Figure 2.**
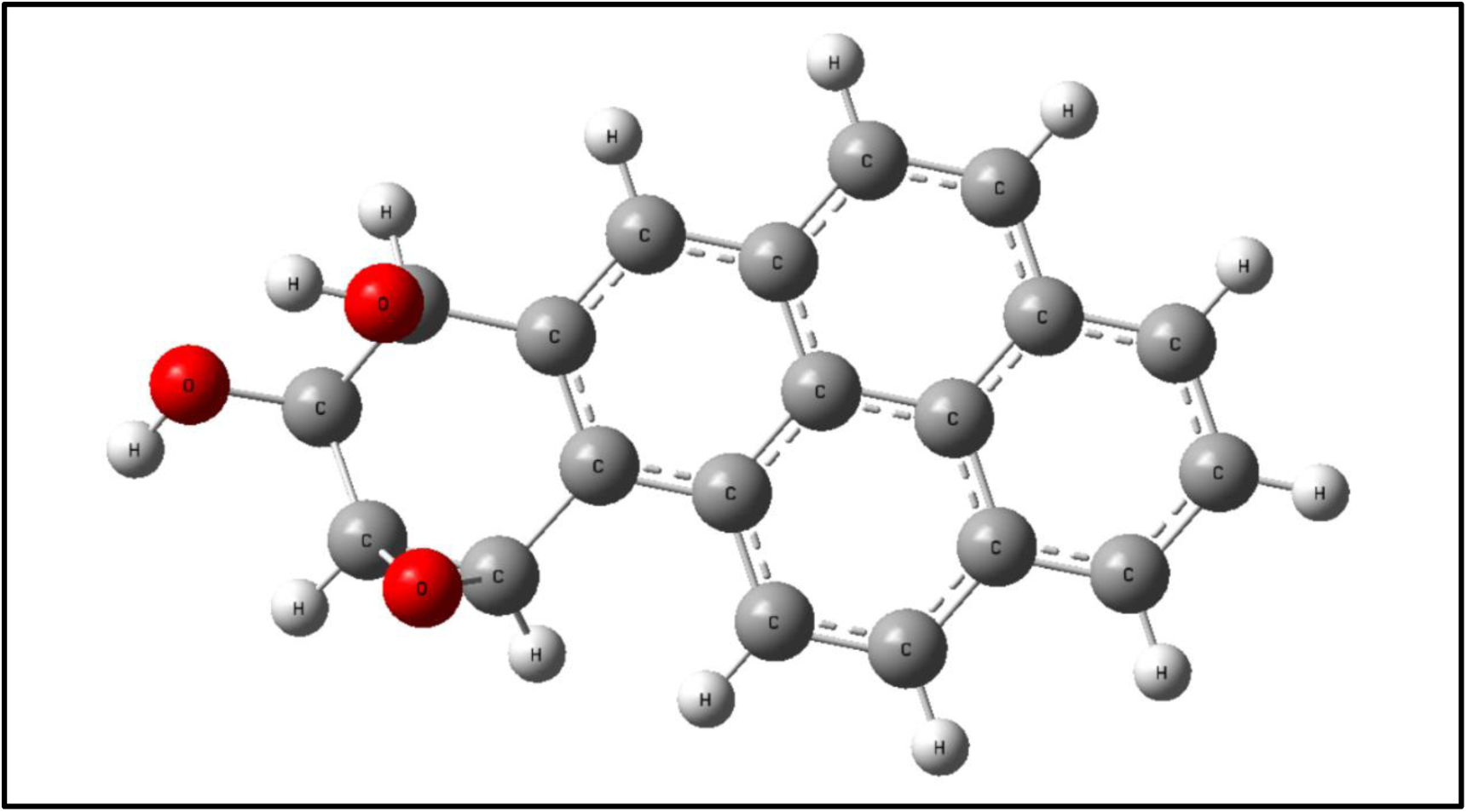
3D ground state configuration of BPDE after optimization

The detail information for bond length, bond angle and dihedral angle for each atom of BPDE molecule including energy minimization and RMS gradient normalization with optimization step number has been shared in supplementary data. After optimisation, the bond lengths in the BPDE molecular structure show no change while bond angle shows slight modifications Figure 2.

### 3.2. Interaction of BPDE and CDK

In Figures 3 (Ia) & (Ib), the docking interactions of BPDE with CDK1 are shown with detailed analyses of hydrophobic and hydrogen bond interactions. Hydrophobic interactions are observed with residues Lys9(E), Leu31(D), Asn45(H), Lys30(D), and Pro33(D) while hydrogen bonds stabilizing the complex are formed between Thr35(D) (3.0 Å), Lys34(D) (2.80 Å), and Tyr19(E) (3.09 Å and 2.92 Å). Van der Waals interactions are seen with His35(D), Asp45(H), and Lys30(D), while pi-alkyl interactions occur between Lys9(E), Lys9(D), and Pro33(D). The 3D representation in Figure 3 (Ic) illustrates the spatial arrangement of these interacting residues. In Figures 3 (IIa) & (IIb), BPDE has hydrogen bonds with Glu12(A) (3.18 Å) and Lys33(A) (3.26 Å), as well as hydrophobic contacts from Leu83(A), Leu134(A), Ala31(A), and others. An interaction map also illustrates hydrogen bonds, carbon-hydrogen bonds with Asp145(A) and Gly13(A), and pi-sigma and pi-alkyl interactions with Leu134(A), Glu32(A), and Lys33(A). The spatial orientation of BPDE within the active site is presented in Figure 3 (IIc), in which the hydrogen bonds and close positioning of BPDE are highlighted. Further, to Figures 3 (IIIa) & (IIIb), the details of BPDE-CDK4 interaction are given where hydrogen bonds form between BPDE and Val247(C) at distances of 2.94 Å and 3.10 Å, respectively. Hydrophobic interactions involve Glu240(C), Pro244(C), Trp243(C), and others. The interaction map shows conventional hydrogen bonds with Val247(C), van der Waals interactions with several residues, and pi-donor hydrogen bonding with Arg251(C). Figure 3 (IIIc) is a 3D view of BPDE bound to the CDK4 active site. Finally, in Figures 3 (IVa) & (IVb), BPDE interacts with CDK6 through hydrogen bonds with Val247(A) (2.59 Å and 2.84 Å) and hydrophobic interactions with Glu240(A), Trp243(A), and other residues. The interaction map shows hydrogen bonds, van der Waals interaction with multiple residues, and pi-alkyl interaction with Trp243(A) and Pro244(A), and finally, pi-cation interaction with Arg245(A).

**Figure 3.**
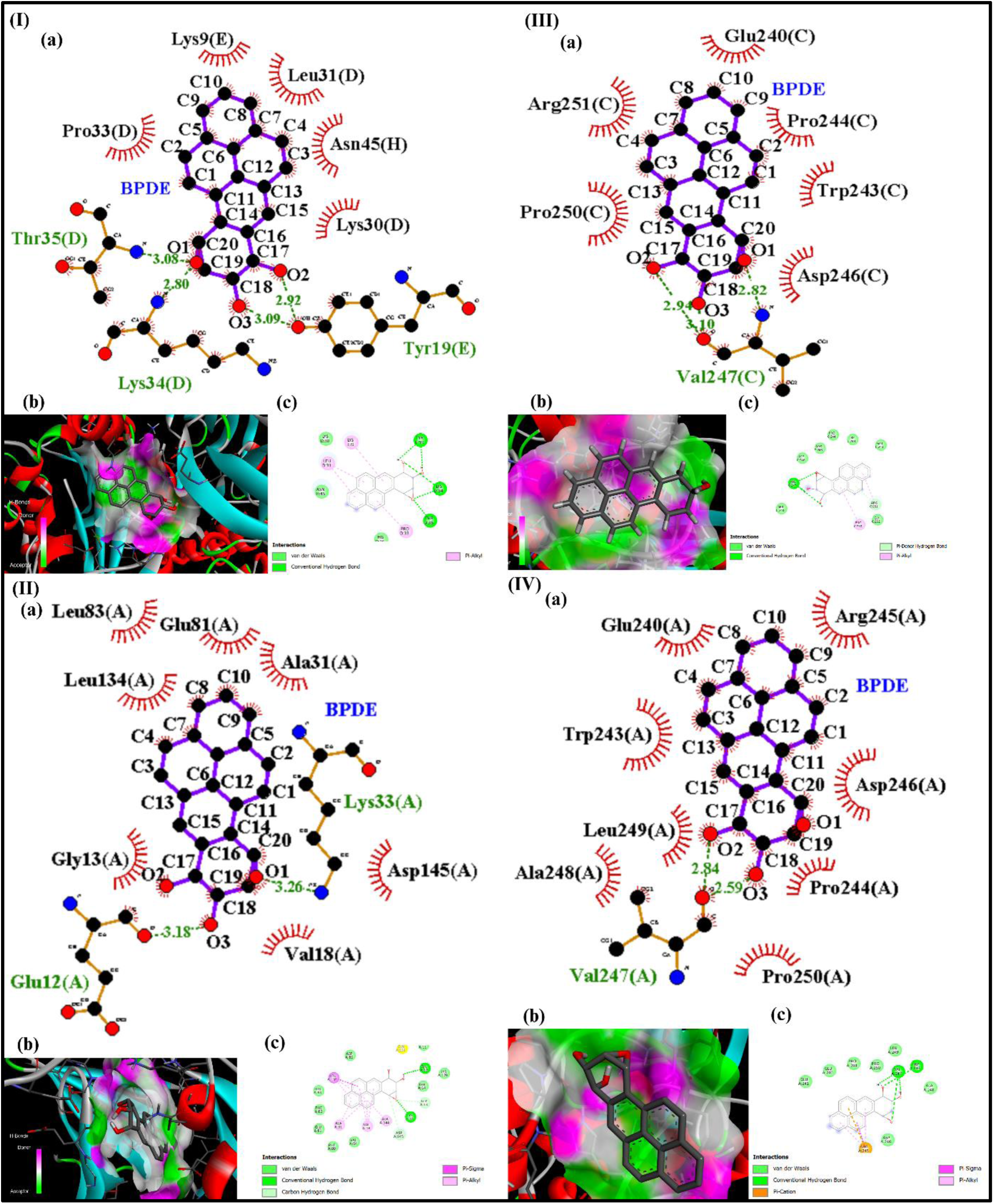
(1) Molecular docking studies of BPDE with CDK1, (a) LigPlot diagram illustrating non-bonded interactions of BPDE-CDK1 complex, (b) 2D representation of key interactions of BPDE-CDK1 complex, (c) 3D Discovery Studio visualization of the binding pose, emphasizing key residues involved in the interaction and the overall binding mode; (2) Molecular docking studies of BPDE with CDK2, (a) LigPlot diagram illustrating non-bonded interactions of BPDE-CDK2 complex, (b) 2D representation of key interactions of BPDE-CDK2 complex, (c) 3D Discovery Studio visualization of the binding pose, emphasizing key residues involved in the interaction and the overall binding mode; (3) Molecular docking studies of BPDE with CDK4, (a) LigPlot diagram illustrating non-bonded interactions of BPDE- CDK4 complex, (b) 2D representation of key interactions of BPDE- CDK1 complex, (c) 3D Discovery Studio visualization of the binding pose, emphasizing key residues involved in the interaction and the overall binding mode; (4) Molecular docking studies of BPDE with CDK6. (a) LigPlot diagram illustrating non-bonded interactions of BPDE- CDK1 complex, (b) 2D representation of key interactions of BPDE- CDK1 complex, (c) 3D Discovery Studio visualization of the binding pose, emphasizing key residues involved in the interaction and the overall binding mode.

**Figure 4.**
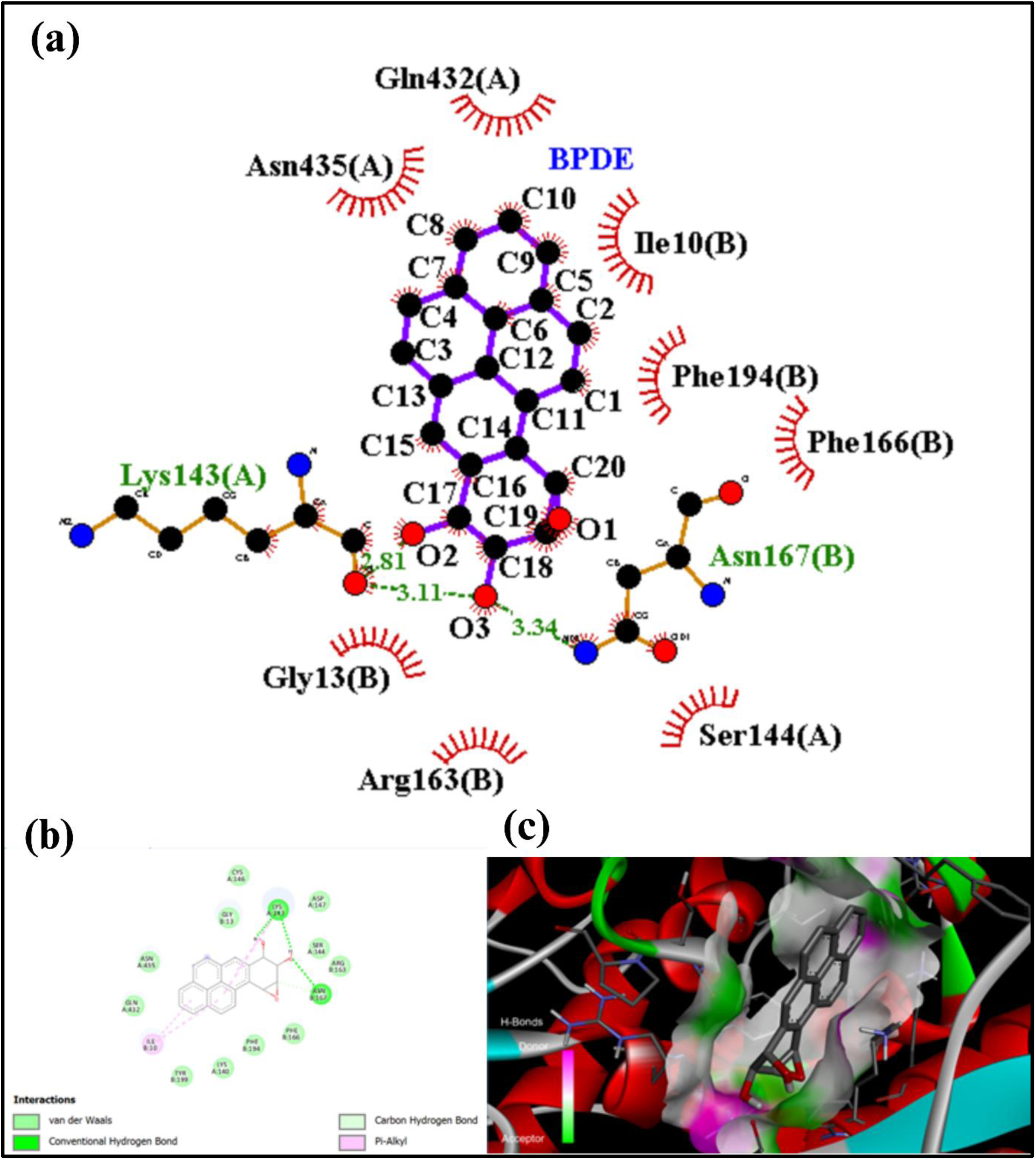
Molecular docking studies of BPDE with Pol η, (a) LigPlot diagram illustrating non-bonded interactions of BPDE-Pol η complex, (b) 2D representation of key interactions of BPDE-Pol η complex, (c) 3D Discovery Studio visualization of the binding pose, emphasizing key residues involved in the interaction and the overall binding mode.

The 3D visualization in Figure 3 (IVc) highlights the binding of BPDE within the CDK6 active site, with clear representations of hydrogen bonds and pocket surface.

### 3.3. Interaction of BPDE and Pol η

The docking interaction results of BPDE with the Pol η are illustrated in Figure 7 (a), the 2D interaction diagram shows that BPDE interacts through hydrogen bonds with residues Lys143(A) at a distance of 3.11 Å and Asn167(B) at a distance of 3.34 Å. Hydrophobic interactions are seen with residues Gln432(A), Asn435(A), Gly13(B), Arg163(B), Ile10(B), Ser144(A), Phe166(B), and Phe194(B). These interactions stabilize BPDE in the protein pocket. In Figure 7 (b), the interaction diagram shows a variety of types of bondings, conventional hydrogen bonds are seen between BPDE and residues Lys143(A) and Asn167(B), with bond length of about 3.11 Å and 3.34 Å, respectively. Pi-alkyl interaction is also formed between aromatic rings of BPDE and the nonpolar side chain of Phe194(B) and Ile10(B), which contributes to the hydrophobic stabilization. There are also weaker van der Waals interactions with surrounding residues, including Gln432(A), Asn435(A), Gly13(B), Arg163(B), Ser144(A), Phe166(B), and Phe194(B). Altogether, hydrogen bonds, Pi-alkyl interactions, and van der Waals forces play a crucial role in the stabilization of BPDE binding into the protein’s active site. Figure 7 (c) shows the binding site as a 3D visualization in which hydrogen bonds are drawn as green dashed lines between BPDE and Pol η. The surface of the binding pocket, with its hydrophobic interactions and Pi-alkyl contacts, again drives home the stable placement of BPDE in the protein.

### 3.4. Interaction of BPDE and HMGB1

In Figure 5 (a), the 2D interaction diagram illustrates that BPDE interacts through hydrogen bonds with residues Glu49(A) at a distance of 2.90 Å (O3) and 2.74 Å (O2), Trp53(A) with O1 at 2.71 Å, and O2 at 3.01 Å. These hydrogen bonds are vital for stabilizing the BPDE molecule within the binding site. Hydrophobic interactions are present with residues Arg26(A), Thr41(A), Val19(A), Asp22(A), Tyr23(A), Met45(A), Met52(A), and Trp15(A), which collectively ensure the stable positioning of BPDE within the HMGB1 protein pocket. In Figure 5 (b), the interaction diagram categorizes the bonding types in greater detail. Conventional hydrogen bonds (depicted by green lines) exist between BPDE and residues Glu49(A) and Trp53(A), ensuring polar stabilization. Pi-alkyl interactions (pink lines) occur between BPDE’s aromatic rings and residues Met45(A). Pi-donor hydrogen bonds are also noted, highlighting unique hydrogen bonding involving π electron systems. Lastly, weak van der Waals interactions with residues Arg26(A), Thr41(A), Val19(A), Asp22(A), Tyr23(A), Met52(A), and Trp15(A) provide further non-covalent stabilization, strengthening BPDE’s binding affinity. Figure 5 (c) offers a 3D representation of the binding pocket, showing the hydrogen bonds between BPDE and residues of HMGB1, represented as green dashed lines on the surface of the binding site. This 3D visualization highlights the spatial arrangement of BPDE within the pocket and underscores the significance of polar and non-polar interactions in stabilizing the molecule within the HMGB1 protein.

**Figure 5.**
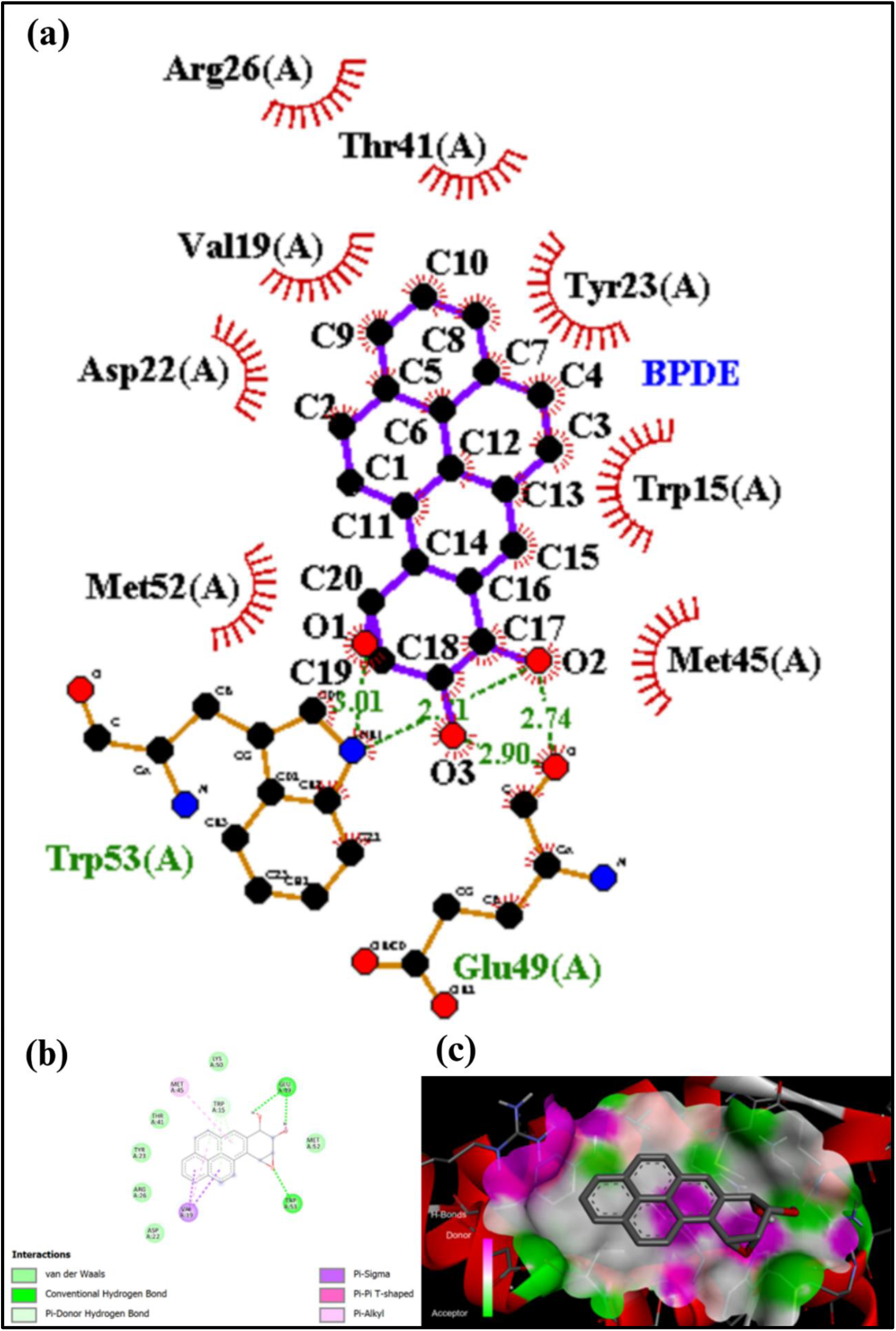
Molecular docking studies of BPDE with HMGB1, (a) LigPlot diagram illustrating non-bonded interactions of BPDE-HMGB1 complex, (b) 2D representation of key interactionsof BPDE-HMGB1 complex, (c) 3D Discovery Studio visualization of the binding pose, emphasizing key residues involved in the interaction and the overall binding mode.

### 3.5. Interaction of BPDE and LOX

Figure 6 (a), the type of the interaction between BPDE and LOX are described in details. The residues of the interacting protein BPDE form two classical hydrogen bonds with Gln434(A) and Gln437(B), with a bond distance of 2.26 Å and 2.54 Å. Further, the phenyl rings of BPDE also donate to the formation of Pi-cation interactions with the help yardstick Arg438(A), Arg438(B) and Lys441(A). Other residues implicated in van der Waals interactions are Pro149(A), Gly150(A), Asp290(B), Gly291(B), and Gln437(A). A Pi-alkyl interaction is also observed with Arg438(A), supporting role of hydrophobic interactions in the stabilization of BPDE at the binding site. Figure 6 (b) the interaction map gives the interaction of BPDE and LOX in a more simplified form in the form of a diagram. Other hydrogen part includes Gln434(A) and Gln437(B) and they are normal hydrogen bonds which are shown as green dashed line and stabilize BPDE. The Pi-cation and Pi-alkyl interactions (purple and orange dashed lines) are with Arg438(A), Arg438(B), and Lys441(A) and the aromatic ring of BPDE. Cis-pi stacking interactions(violet line) are noted with NH2 groups of Pro149(A) and Gly150(A), COO groups of ASP290(B) and GLY291(B), and backbone amide group of Gln437(A) and also Van der waal interactions (green bubbles) are seen with Pro149(A), Gly150(A), Asp290(B), Gly291(B) and Gln437(A). As seen from this map, several types of non-coalvalent interactions are involved in the docking stability of BPDE. Figure 6 (c) the 3D interaction visualization demonstrates the position of BPDE in the LOX binding pocket. The chiral centers of BPDE form Pi-cation and Pi-alkyl interactions with the positively charged residues Arg438(A), Arg438(B), and Lys441(A) of the aromatic rings. Hydrogen bonds with Gln434(A) and Gln 437(B) are also observed firmly attaching BPDE at the pocket. The residues Pro149, Gly150, Asp290 and Gly291 are observed to form the contacts with BPDE by means of van der Waals’ interactions making the complex more stable. This visualization substantiates the synergistic interaction of the H-bonds, Pi-interaction and van der Waal forces in stabilizing binding of BPDE to LOX.

**Figure 6.**
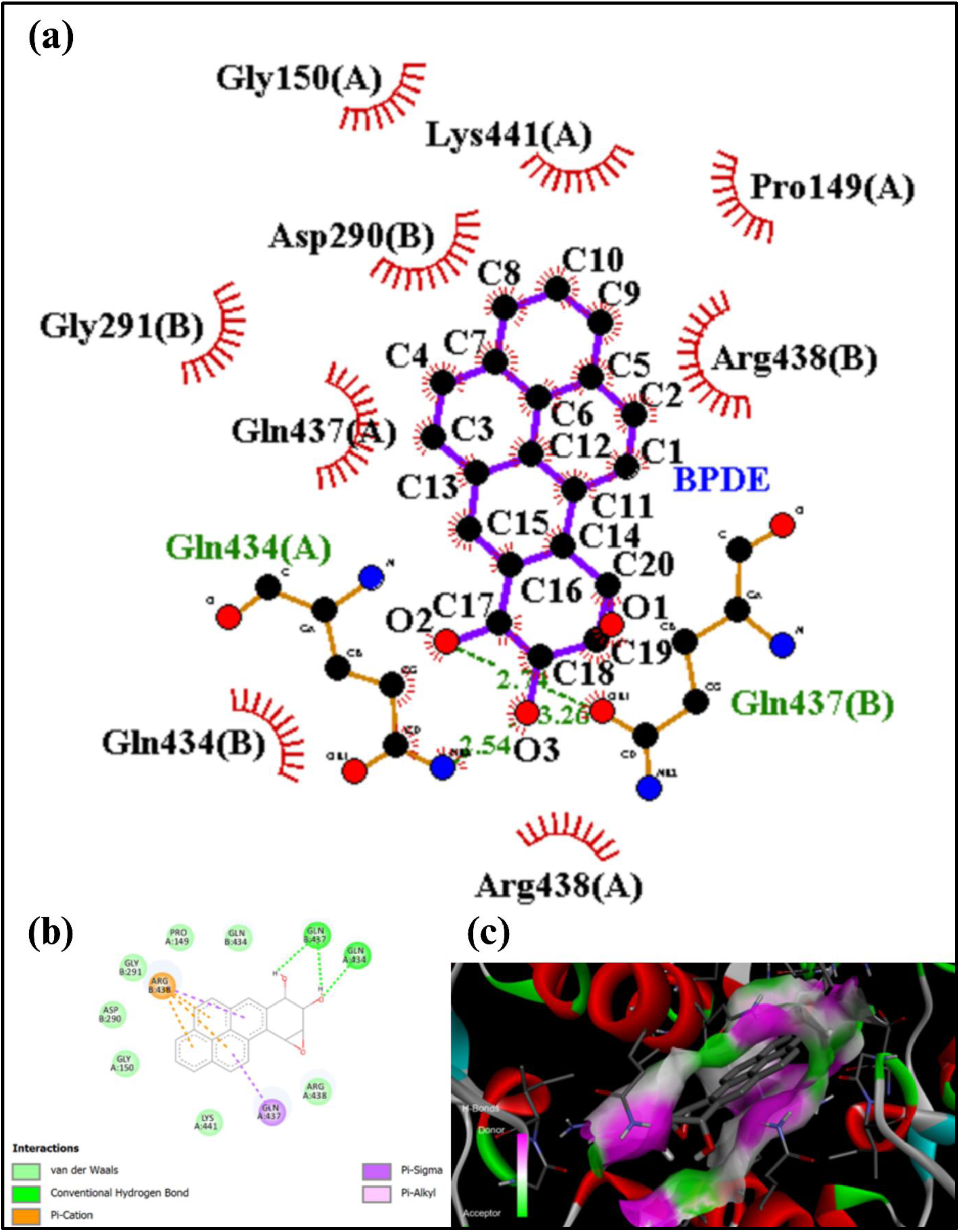
Molecular docking studies of BPDE with LOX, (a) LigPlot diagram illustrating non-bonded interactions of BPDE-LOX complex, (b) 2D representation of key interactions of BPDE-LOX complex, (c) 3D Discovery Studio visualization of the binding pose, emphasizing key residues involved in the interaction and the overall binding mode.

### 3.6. Interaction of BPDE and p53

Figure 7 (a) in the 2D interaction diagram, BPDE interacts with p53 by hydrogen bonds, carbon-hydrogen bonds, Pi-interactions, and van der Waals forces. Conventional hydrogen bonds are formed by BPDE with Tyr107(A), where oxygen atoms O3 of BPDE are interacting at bond lengths of 3.03 Å and 3.20 Å, thus stabilizing it in the binding site. A carbon-hydrogen bond is also found with Asn239(C). The aromatic rings of BPDE engage in Pi-interactions: amide-Pi stacking with residues Pro152(A) and Pro153(A), and Pi-alkyl interactions with residues such as Leu137(C) and Ala276(C). The van der Waals forces also play a role in stabilization for several residues, including Gln136(C), Lys139(C), Thr150(A), Cys275(C), and Leu137(C). Figure 7 (b) the interaction map schematically shows the binding of BPDE to p53. The conventional hydrogen bonds are represented by green dashed lines, which show interactions with Tyr107(A). A carbon hydrogen bond is seen with Asn239(C), while Pi-interactions are marked with purple and orange dashed lines, showing amide-Pi stacking and Pi-alkyl interactions with Pro152(A), Pro153(A), Leu137(C), and Ala276(C). The van der Waals interactions, depicted as green bubbles, include residues such as Gln136(C), Lys139(C), Thr150(A), Cys275(C), and Leu137(C). These collectively stabilize BPDE within the p53 binding pocket and highlight the importance of aromatic and hydrophobic contacts. Figure 7 (c) the spatial orientation of BPDE within the p53 binding pocket is visualized through the 3D interaction visualization. BPDE’s aromatic rings have established amide-Pi stacking and Pi-alkyl interactions with residues Pro152(A), Pro153(A), Leu137(C), and Ala276(C). Pi-interactions also bring into view the role of hydrophobic as well as aromatic contacts for BPDE’s binding. The conventional hydrogen bonds with Tyr107(A) are visible, which anchor BPDE through polar interactions. Residues such as Asn239(C), Gln136(C), Lys139(C), Thr150(A), and Cys275(C) contribute through van der Waals forces, further stabilizing BPDE in the pocket. The combination of hydrogen bonding, Pi-interactions, and van der Waals forces ensures a strong and cooperative binding of BPDE to p53.

### 3.7. Interaction of BPDE and EGFR

Figure 8 (a) the 2D interaction diagram of BPDE docked with EGFR shows that there are a number of stabilizing interactions, including hydrogen bonds, Pi-interactions, and van der Waals forces. A conventional hydrogen bond is formed with Gly983(A), with the O3 atom of BPDE interacting at a bond length of 2.76 Å. Van der Waals forces have been contributed by several residues, such as Lys806(A), Gln982(A), Glu985(A), Asp984(A), and Asp807(A), for increasing the binding stability. In addition, amide-Pi stacking and Pi-alkyl interactions take place between the aromatic core of BPDE and residues like Ile809(A), Ile981(A), and Glu985(A). These interactions firmly anchor BPDE within the EGFR binding pocket, and both aromatic and polar residues are critical in this regard. Figure 8 (b) the interaction map depicts the detailed molecular interaction of BPDE with EGFR. The conventional hydrogen bond with Gly983(A) is shown as green dashed lines, which anchors BPDE. Pi-alkyl interactions are shown in purple, which include hydrophobic residues such as Ile809(A) and Ile981(A). The amide-Pi stacked interactions are seen with residues like Glu985(A). In addition, residues like Asp807(A), Lys806(A), and Asn808(A) engage in van der Waals interactions, represented as green bubbles, contributing to the overall binding stability of BPDE within EGFR. Figure 8 (c) the 3D view clearly indicates the spatial positioning of BPDE within the EGFR binding pocket using Discovery Studio.

**Figure 7.**
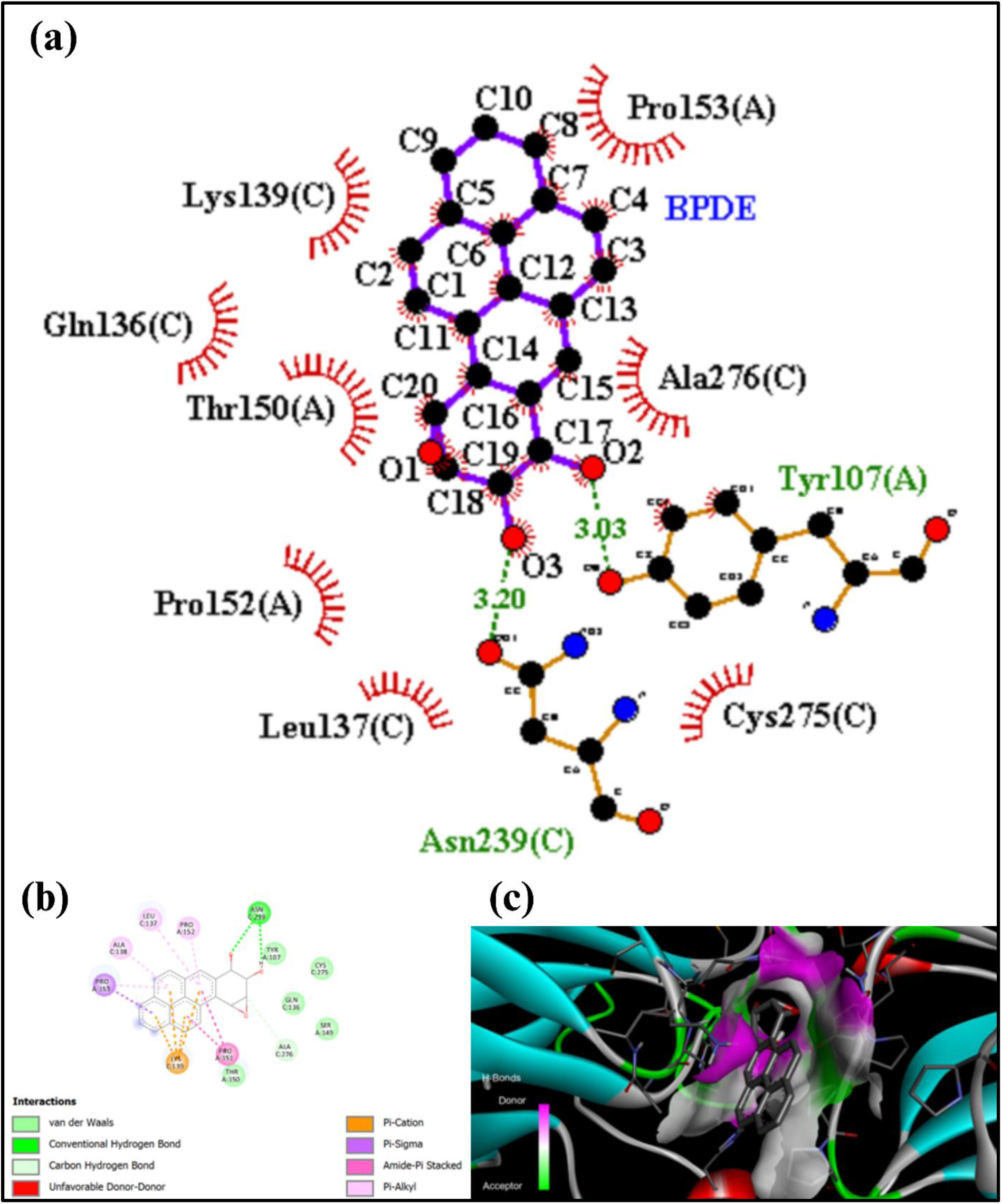
Molecular docking studies of BPDE with p53, (a) LigPlot diagram illustrating non- bonded interactions of BPDE-p53 complex, (b) 2D representation of key interactions of BPDE- p53 complex, (c) 3D Discovery Studio visualization of the binding pose, emphasizing key residues involved in the interaction and the overall binding mode.

**Figure 8.**
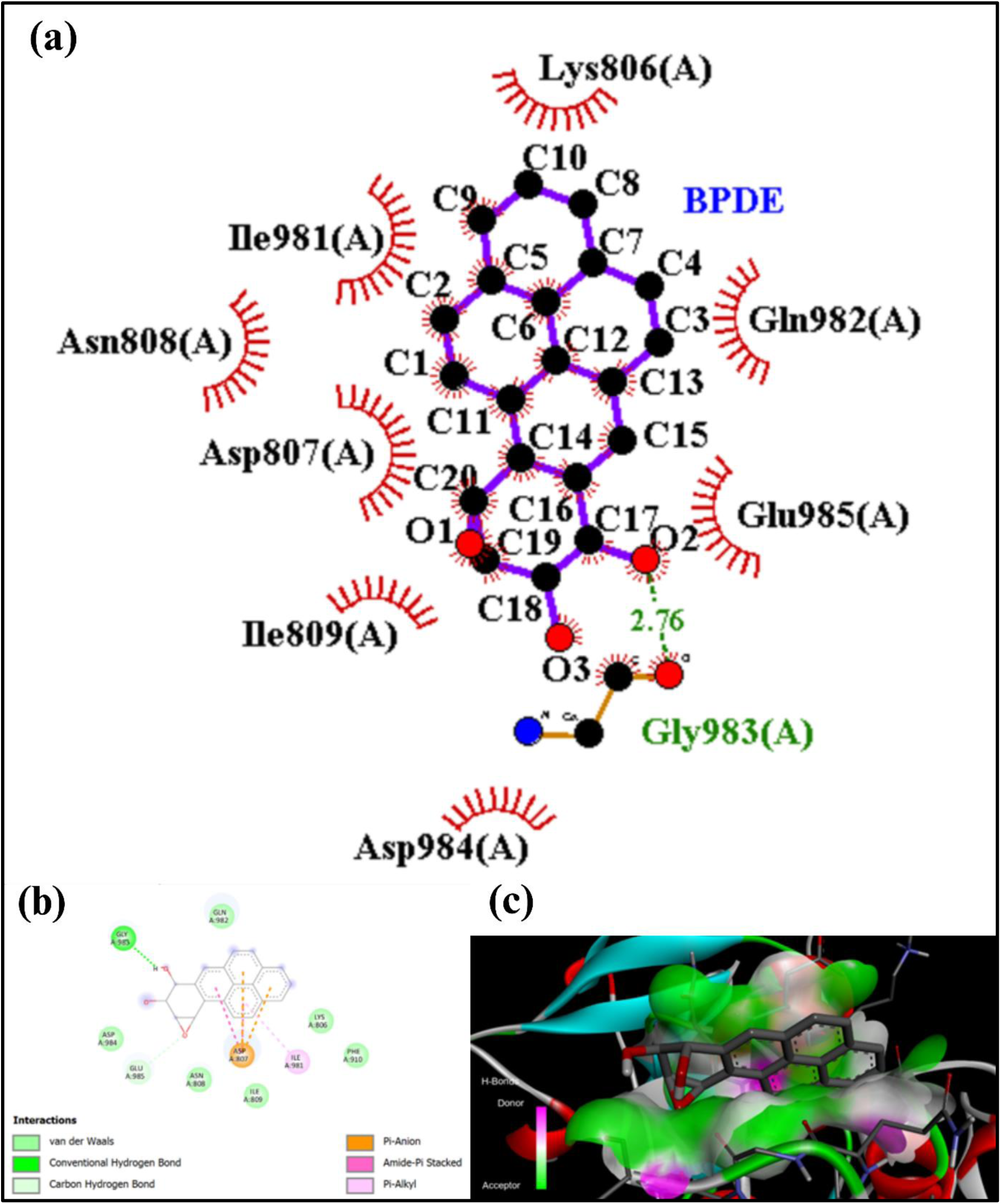
Molecular docking studies of BPDE with EGFR, (a) LigPlot diagram illustrating nonbonded interactions of BPDE-EGFR complex, (b) 2D representation of key interactions of BPDE-EGFR complex, (c) 3D Discovery Studio visualization of the binding pose, emphasizing key residues involved in the interaction and the overall binding mode.

### 3.8. Interaction of BPDE and NF-κB

Figure 9 (a) the 2D diagram shows the docking interaction of BPDE with NF-κB. BPDE forms two conventional hydrogen bonds with key residues Leu280(F) and Lys221(A). The first bond involves the O3 atom of BPDE interacting with Leu280(F) at a bond distance of 3.32 Å, while the second interaction with Lys221(A) occurs at 3.06 Å. In addition, van der Waals forces play a critical role in stabilizing the interaction with residues such as His245(A), Gln249(F), Tyr251(F), Pro281(F), and Arg246(A). The hydrophobic residues Val244(A) and Val254(B) are also involved; they enhance the stability of BPDE through non-polar interactions. Figure 9 (b) the interaction map gives a vivid picture of the molecular interactions. Hydrogen bonds between BPDE and Leu280(F) and Lys221(A) are very crucial for anchoring the molecule within the binding site. Additional Pi-alkyl interactions were also observed with residues such as Tyr251(F) and Pro281(F), which are essential for hydrophobic stabilization of BPDE. Pi-cation interactions were noted, which further strengthens the docking position. Van der Waals forces, depicted as green bubbles, are ubiquitous and involve residues such as Arg246(A), Val244(A), and His245(A), thus ensuring overall stability of the BPDE-EGFR/NF-κB complex. Figure 9 (c) the 3D visualization brings to light the spatial distribution of BPDE within the NF-κB binding pocket.

**Figure 9.**
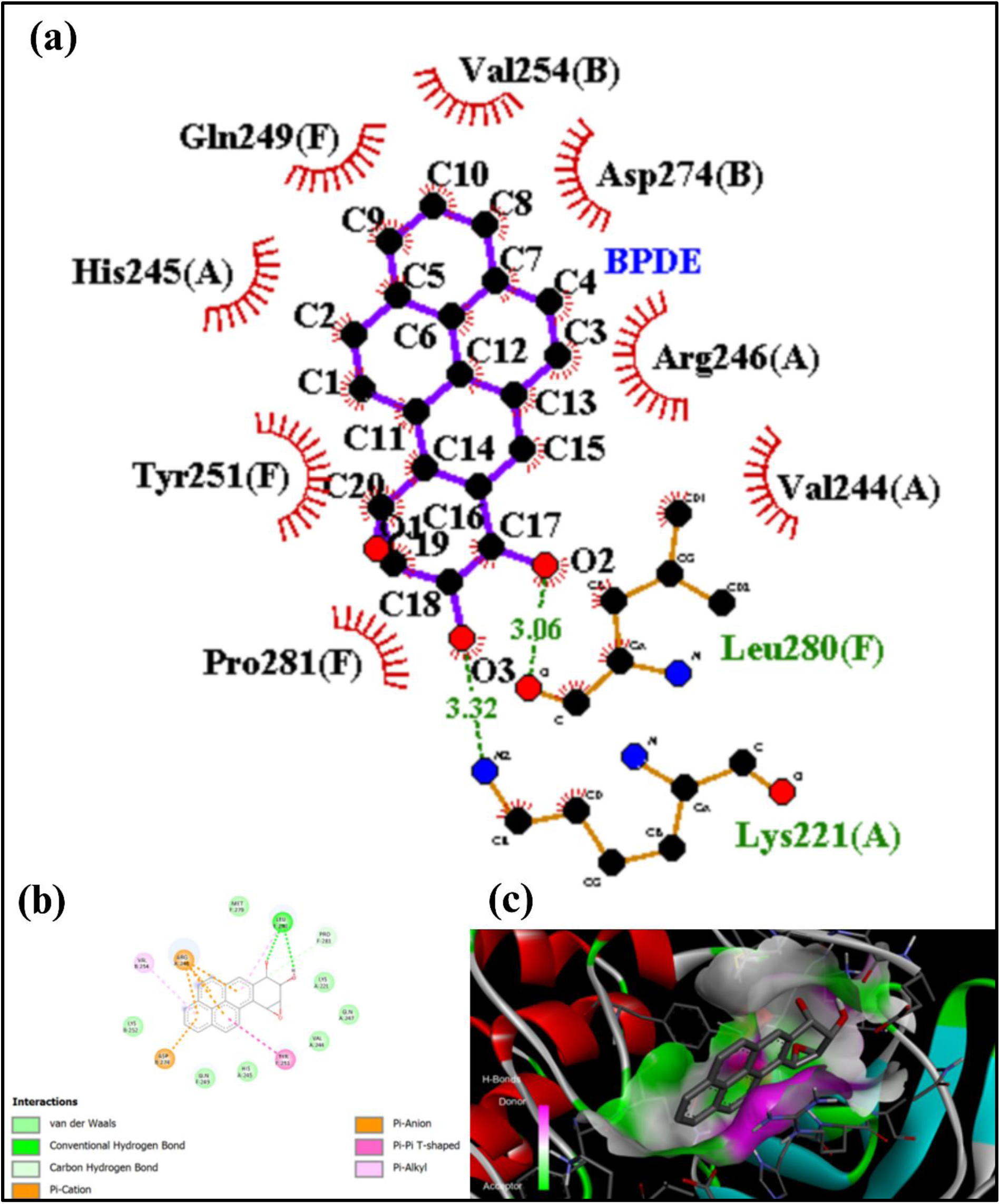
Molecular docking studies of BPDE with NF-κB, (a) LigPlot diagram illustrating non-bonded interactions of BPDE-NF-κB complex, (b) 2D representation of key interactions of BPDE-NF-κB complex, (c) 3D Discovery Studio visualization of the binding pose, emphasizing key residues involved in the interaction and the overall binding mode.

The current study applied molecular docking to analyze the interaction of BPDE with significant proteins implicated in the carcinogenic process, such as NF-κB, MAPK, and PI3K/Akt signaling pathways. Figure 10 analyses provide insights into molecular details that are often difficult to achieve through strictly experimental procedures. The docking results revealed a moderate interaction for the BPDE-NF-κB complex with a binding energy of -6.11 kcal/mol, when compared to other cascadic proteins (Figure 13). The docking conformation of BPDE with NF-κB, as illustrated by molecular PYMOL, clearly reveals the binding of the ligand at the active site of the protein. It involves both hydrogen bonds and hydrophobic interaction between the ligand and protein. Most importantly, this interaction also involves conventional hydrogen bonds by AA presents hydrogen bonds and more enhanced hydrophobic interactions along with AA. The affinities of BPDE binding towards other crucial proteins were apparently higher than that for the BPDE-NF-κB complex, revealing even better targets in most instances.

**Figure 10.**
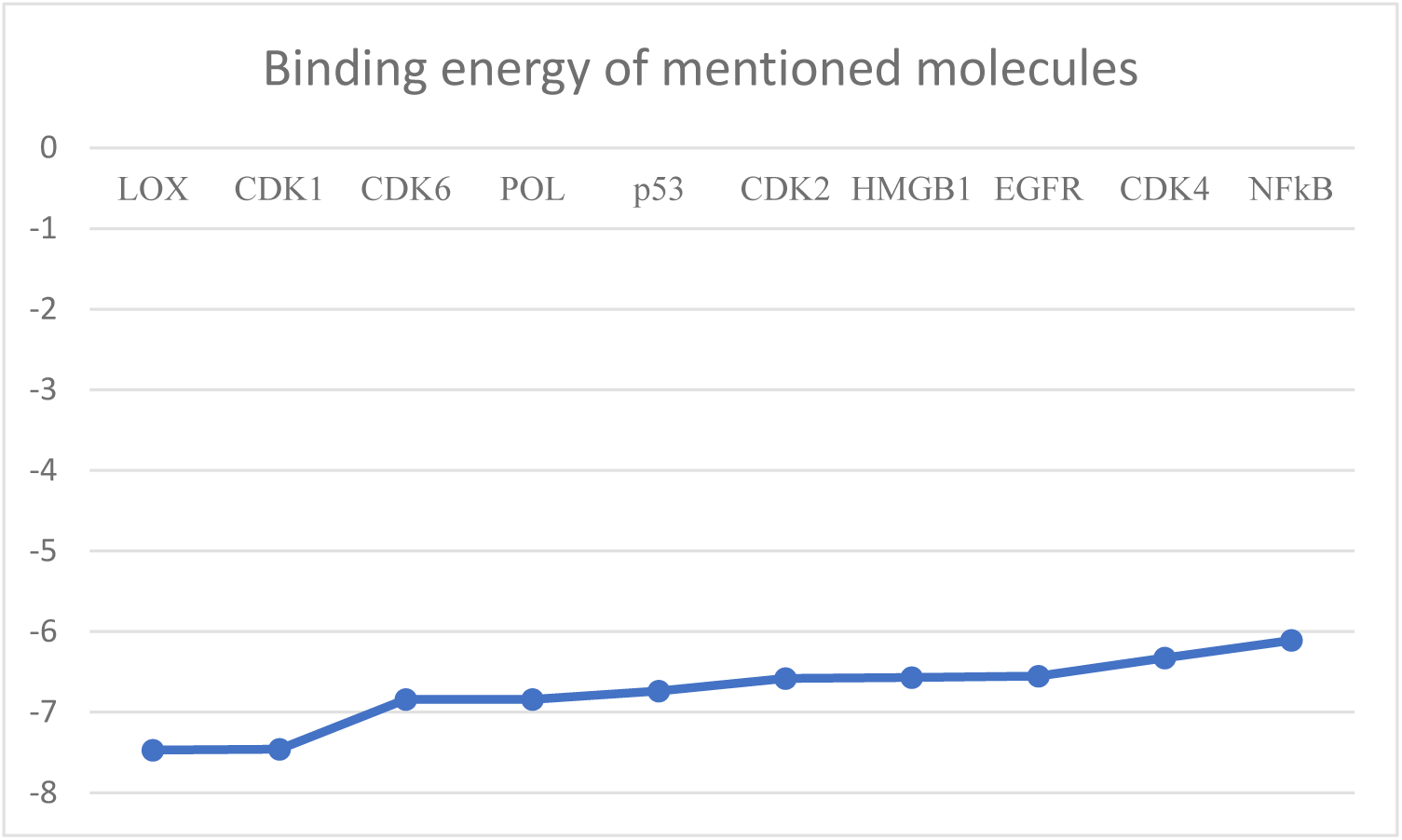
Graph showing binding energy values of BPDE interaction with different cascading pathway receptor molecules.

**Figure 11.**
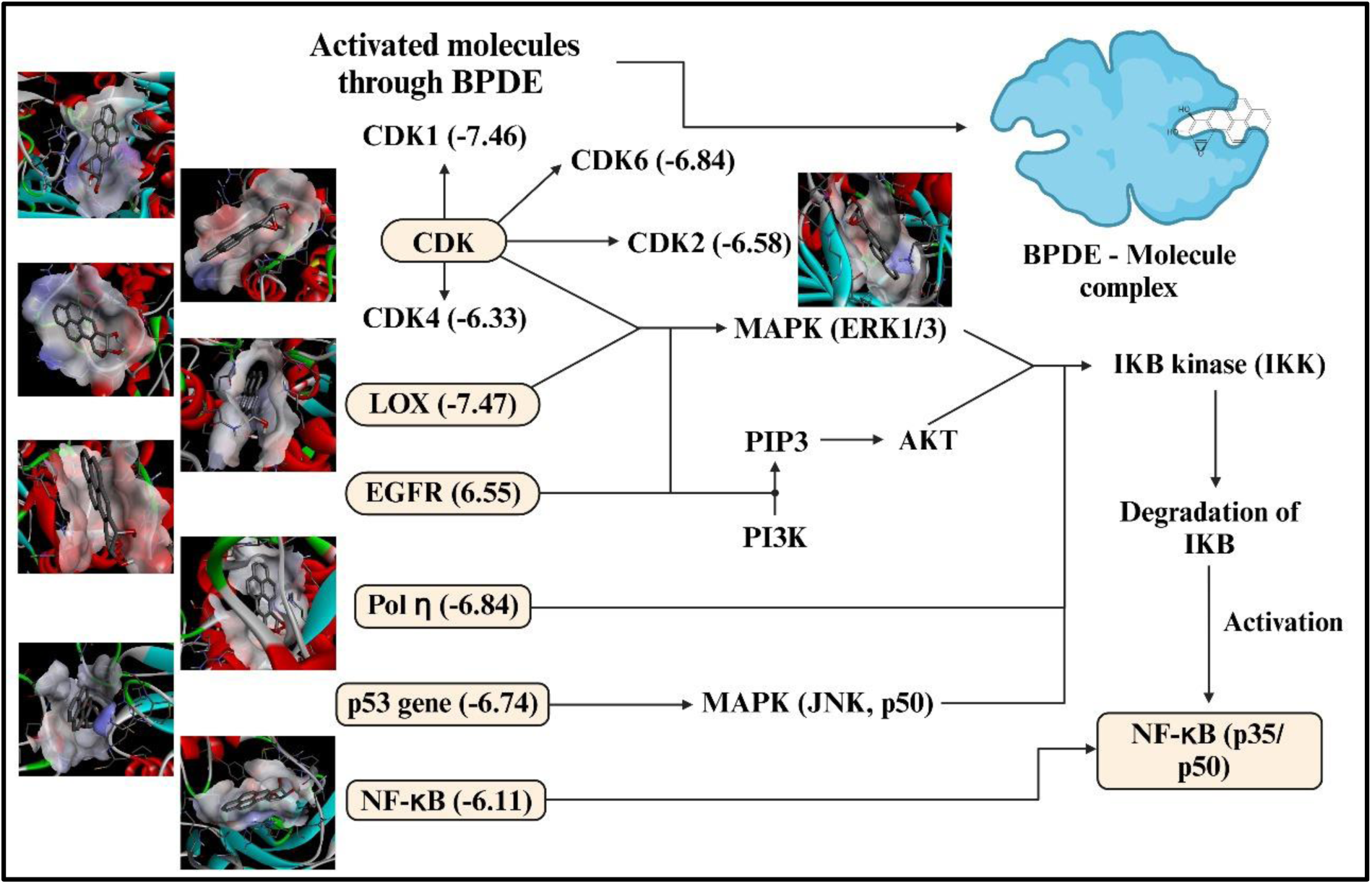
The proposed mechanism of cell signaling, cascading from BPDE through different molecules till NF-κB; Mentioned binding energy is in (kcal/mol).

Binding energy with maximum value of -7.47 kcal/mol with AA comprising the major interactions of BPDE towards LOX. Besides, a binding energy of -6.84 kcal/mol was obtained in the case of the BPDE-Pol η complex where key residues such as AA were engaged in hydrogen bonding interactions. The interaction of BPDE with CDK6 also appeared to be strong with a binding energy of -6.84 kcal/mol wherein the role of AA residues could be visualized during interaction. A binding energy of -7.46 kcal/mol was obtained for the BPDE-CDK1 complex in which residues of AA had contributed. Similar outcomes have been seen in case of BPDE-CDK2 and BPDE-CDK4 with respective binding energies of -6.58 kcal/mol and -6.33 kcal/mol. The BPDE-EGFR complex fell into the category of moderate interactions with a binding energy of -6.55 kcal/mol, mainly through hydrophobic and polar interactions involving AA. BPDE-p53 and BPDE-HMGB1 complexes were calculated to have binding energies of -6.74 kcal/mol and -6.57 kcal/mol, respectively, where residues like Cysteine 242 (p53) and Tryptophan 42 (HMGB1) appeared critical.

These data suggest that although BPDE has a moderate affinity for NF-κB, it exhibits stronger binding affinities for proteins associated with carcinogenic signaling pathways. Therefore, BPDE appears to interact with multiple crucial proteins that support its multiple etiologic contributions in carcinogenesis. Molecular mechanisms of the interaction between BPDE and target proteins should be focused. Knowledge of these interactions at the more profound level may lead to developing specific inhibitors for high affinity targets like 5-Lipoxygenase (LOX) and CDK6 as some future therapy in cancer treatment. Testing combined therapeutic strategies in preclinical and clinical models may also give insight into the efficacy of targeting multiple pathways.

This study’s relevance extends into the arena of public health, mainly into that of cancer prevention and treatment. The determination of the biomarkers of BPDE exposure will provide an early and straightforward approach to monitor and detect carcinogenicity among susceptible populations. The understanding of signaling pathways of BPDE activation may enable public health interventions aimed at reduction in the exposure of susceptible populations to this carcinogen and targeted intervention programs.

The public health context highlights the significance of understanding molecular interactions of carcinogens such as BPDE. Activation of the MAPK and PI3K/Akt signaling pathways, involving proteins such as HMGB1 and CDKs, underlines the complexity of the biology of cancer. In this respect, public health strategies may be enhanced through the incorporation of molecular research findings into programs for the prevention of cancer, especially within communities that have a high level of exposure to environmental carcinogens. With the discovery of signaling pathways as targets for inhibition, new drugs may be developed to provide a much better outcome for patients with cancer. By identification of the molecular mechanisms of carcinogenesis, healthcare professionals may design interventions targeting both the manifestations of cancer and the driving mechanisms behind the cancerous tumour.

## 4. Proposed future BPDE-induced mechanism

BPDE exhibits varying binding affinities with a range of critical molecules, influencing diverse cellular pathways. Among these, CDK1 demonstrates the strongest interaction (-7.46 kcal/mol) among the cyclin-dependent kinases, followed by CDK6 (-6.84 kcal/mol), CDK2 (-6.58 kcal/mol), and CDK4 (-6.33 kcal/mol). This hierarchy of binding suggests that BPDE preferentially targets CDK1, likely disrupting cell cycle regulation more significantly through this protein. CDK1’s much higher affinity may have drastically altered effects on cell cycle progression, resulting in enhanced proliferation, while the relatively lower affinities for CDK2, CDK4, and CDK6 may result in secondary progression of checkpoints in a cell cycle.

Interaction to LOX (-7.47 kcal/mol) is the best of its binding among the inflammatory pathway targets because it surpasses the relative affinity seen for NF-κB (-6.11 kcal/mol). This implies that BPDE may significantly induce the production of pro-inflammatory mediators like leukotrienes, thus possibly supporting the tumour microenvironment (TME) and chronic inflammation. The interaction of HMGB1 is likely to mainly result in oxidative stress and inflammatory receptor activation such as RAGE and TLR4. Among the DNA repair and tumour-suppressor proteins, BPDE binds most strongly to POL (-6.84 kcal/mol) and closely to p53 (-6.74 kcal/mol). Interaction with POL suggests a critical disruption of DNA repair mechanisms, whereas binding to p53 likely disrupts its tumour-suppressor functions leading to genomic instability. The latter is inter-related, since inhibition of POL in DNA repair would make worse the mutagenic consequence of p53 dysfunction.

For growth and survival pathways, BPDE’s affinity for EGFR (-6.55 kcal/mol) suggests moderate activation of signaling cascades like PI3K/AKT for cell proliferation and resistance to apoptosis. Compared to other inflammatory molecules such as LOX and HMGB1, its affinity for EGFR is slightly lower and thus might be more balanced but impactful on cell growth and survival. NF-κB (-6.11 kcal/mol) is the weakest interacting one among the compounds assayed here, but the induced response is still a meaningful promoter for pro-inflammatory cytokines production and anti-apoptotic signaling. Being weaker, NF-κB function may be ancillary as it enhances cumulative inflammation and survival signaling instead of providing a driving force compared with stronger binders such as other molecules. The biological impact of BPDE binding would also depend on factors such as the concentration of BPDE, the cellular context, and the presence of other interacting molecules.

## 5. Future prospect

This study builds foundational knowledge on BPDE-induced carcinogenesis in light of its interaction with NF-κB. The interaction of molecular partners includes BPDE/NF-κB, HMGB1, CDKs, Pol η, 5-Lipoxygenase, p53, and EGFR. Notably, a ligand of BPDE targets include CDK6, Pol η, and 5-Lipoxygenase, which could indicate activation of NF-κB successively through MAPK and PI3K/Akt. This raises a question for detailed research on how HMGB1 affects the activation of NF-κB. The differential binding affinities of BPDE present treatment possibilities. From the available EGFR inhibitors (Cetuximab, Osimertinib, Afatinib, Dacomitinib, Gefitinib, and Erlotinib), the combination of HMGB1, CDKs, or MAPK/PI3K antagonists could improve efficacy and have numerous advances in clinical applications. For example, EGFR inhibitors might be synergistic with MAPK and PI3K inhibitors, as targeting the HMGB1 DNA repair pathway might abrogate genomic instability and tumour development. This research can also help to establish biomarkers for the early diagnosis of BPDE-induced carcinogenicity. Knowledge of the molecular interactions of BPDE with proteins such as CDKs and HMGB1 could point to other biomolecular markers that could aid in cancer diagnostics or determine response to treatment. Further work should focus on elucidating BPDE-protein interactions, developing inhibitors against high-affinity targets, and establishing combinational therapies in both animal tests and human trials. This study will help the researcher to synthesize aptamer against BPDE and that will intern help further to develop the biosensor and aptasensor to monitor the cancer along with personalised medicine ^42–47^.

## 6. Conclusion

This paper reports a comprehensive molecular docking analysis of BPDE with NF-κB and other proteins in cellular signaling pathways. We employed AutoDock 4.2 to identify specific binding interactions, negative binding energy values indicating stable ligand-receptor complexes. The findings suggest that BPDE is involved in the regulation of important pathways such as MAPK and PI3K/Akt, which are involved in oncogenesis. The results indicate that interaction of BPDE with NF-κB may play a critical role in the initiation of inflammatory responses and cancer-related signaling, which would provide a molecular basis for understanding its carcinogenic potential. In addition, binding patterns of BPDE with other proteins, such as CDKs, Pol η, and p53, emphasize its wide impact on genomic stability and cell cycle regulation. Although the computational insights predict quite useful interactions, their biological relevance must be experimentally confirmed through additional experiments. Further in vitro and in vivo assays may help clarify functional implications of BPDE binding and further determine its role in the activation of downstream signaling pathways.

## Abbreviations

1. BPDE - Benzo[a]pyrene Diol Epoxide
2. BaP - Benzo[a]Pyrene
3. NF-κB - Nuclear Factor Kappa B
4. MAPK - Mitogen-Activated Protein Kinases
5. PI3K/Akt - Phosphoinositide 3-Kinase / Akt
6. HMGB1 - High Mobility Group Box 1
7. CDK - Cyclin-Dependent Kinases
8. Pol η - DNA Polymerase Eta
9. LOX - 5-Lipoxygenase
10. EGFR - Epidermal Growth Factor Receptor
11. PDB - Protein Data Bank
12. RMSD - Root Mean Square Deviation
13. IL-6 - Interleukin-6
14. IL-1β - Interleukin-1 Beta
15. IL – Interleukin
16. TNF-α - Tumor Necrosis Factor Alpha
17. RAGE - Receptor for Advanced Glycation End-products
18. TLR4 - Toll-Like Receptor 4
19. VEGF - Vascular Endothelial Growth Factor
20. MM2 - Molecular Mechanics 2
21. DFT - Density Functional Theory
22. RB3LYP - Restricted B3LYP Functional (in computational chemistry)
23. AA - Amino Acids
24. GA - Genetic Algorithm
25. PIP3 - Phosphatidylinositol (3,4,5)-trisphosphate
26. PDK1 - Phosphoinositide-Dependent Kinase-1
27. mTORC1 - Mechanistic Target of Rapamycin Complex 2
28. mTOR - Mechanistic Target of Rapamycin
29. IKK - IκB Kinase
30. IκB - Inhibitor of κB
31. CpG - Cytosine and Guanine separated by a phosphate group
32. CpG Islands - Regions rich in cytosine-guanine sequences
33. miRNAs – MicroRNAs
34. DNA - Deoxyribonucleic Acid
35. RNA - Ribonucleic Acid
36. H-bond / H-bonds - Hydrogen Bond(s)
37. RMS Gradient - Root Mean Square Gradient
38. CYP1A1 - Cytochrome P450 Family 1 Subfamily A Member 1
39. CYP1B1 - Cytochrome P450 Family 1 Subfamily B Member 1
40. CYP - Cytochrome P450
41. ASK1 - Apoptosis Signal-Regulating Kinase 1
42. AP-1 - Activator Protein 1
43. PKB/Akt / PKB - Protein Kinase B (another name for Akt)
44. GTP - Guanosine Triphosphate
45. GTPase - Guanosine Triphosphatase
46. PDB ID - Protein Data Bank Identifier
47. TGF-β - Transforming Growth Factor Beta
48. ERK / ERK1/3 - Extracellular Signal-Regulated Kinase(s)
49. JNK - c-Jun N-terminal Kinase
50. MAPKKK - Mitogen-Activated Protein Kinase Kinase Kinase
51. PAH - Polycyclic Aromatic Hydrocarbon
52. ROS - Reactive Oxygen Species
53. RTK - Receptor Tyrosine Kinase
54. TME - Tumor Microenvironment
55. FAD - Flavin Adenine Dinucleotide
56. PHE - Phenylalanine (amino acid)
57. GLY - Glycine (amino acid)
58. SER - Serine (amino acid)
59. ARG - Arginine (amino acid)
60. THR - Threonine (amino acid)
61. ASN - Asparagine (amino acid)
62. VAL - Valine (amino acid)
63. ASP - Aspartic Acid (amino acid)
64. LEU - Leucine (amino acid)
65. ALA - Alanine (amino acid)
66. TYR - Tyrosine (amino acid)
67. CYS - Cysteine (amino acid)
68. LYS - Lysine (amino acid)
69. PRO - Proline (amino acid)
70. MET - Methionine (amino acid)
71. Bcl-2 - B-cell Lymphoma 2
72. Bcl-xL - B-cell Lymphoma-extra Large
73. TAMPs - Tumor-Associated Molecular Patterns
74. RMS - Root Mean Square
75. Pymol - Molecular Visualization Tool

## Institutional Review Board Statement

Not applicable

## Informed Consent Statement

Not applicable.

## Ethics approval and consent to participate

Not applicable

## Consent for publication

Yes

## Availability of data and material

Will be made available upon request

## Author Contribution

KVK and AKV performed the experiment, analysis, graphical designing and wrote the manuscript. AKV led the development of methodology for the experiment, data extraction, study quality assessment, conceptualization, study identification, analysis, manuscript writing and editing, and overall supervision. KVK edited the whole draft and did the referencing. AKV and AK helped preparation, analysis and editing the draft and reviewed the whole draft, edited the paper for final version with crucial feedback.

## Acknowledgement

The authors would like to thank the School of Bioengineering and Biosciences, Lovely Professional University, Phagwara, Punjab, 144411 for providing us with all kinds of support.

